# Electrically silent KvS subunits associate with native Kv2 channels in brain and impact diverse properties of channel function

**DOI:** 10.1101/2024.01.25.577135

**Authors:** Michael Ferns, Deborah van der List, Nicholas C. Vierra, Taylor Lacey, Karl Murray, Michael Kirmiz, Robert G. Stewart, Jon T. Sack, James S. Trimmer

## Abstract

Voltage-gated K^+^ channels of the Kv2 family are highly expressed in brain and play dual roles in regulating neuronal excitability and in organizing endoplasmic reticulum - plasma membrane (ER-PM) junctions. Studies in heterologous cells suggest that the two pore-forming alpha subunits Kv2.1 and Kv2.2 assemble with “electrically silent” KvS subunits to form heterotetrameric channels with distinct biophysical properties. Here, using mass spectrometry-based proteomics, we identified five KvS subunits as components of native Kv2.1 channels immunopurified from mouse brain, the most abundant being Kv5.1. We found that Kv5.1 co-immunoprecipitates with Kv2.1 and to a lesser extent with Kv2.2 from brain lysates, and that Kv5.1 protein levels are decreased by 70% in Kv2.1 knockout mice and 95% in Kv2.1/2.2 double knockout mice. Multiplex immunofluorescent labelling of rodent brain sections revealed that in neocortex Kv5.1 immunolabeling is apparent in a large percentage of Kv2.1 and Kv2.2-positive layer 2/3 neurons, and in a smaller percentage of layer 5 and 6 neurons. At the subcellular level, Kv5.1 is co-clustered with Kv2.1 and Kv2.2 at ER-PM junctions in cortical neurons, although clustering of Kv5.1-containing channels is reduced relative to homomeric Kv2 channels. We also found that in heterologous cells coexpression with Kv5.1 reduces the clustering and alters the pharmacological properties of Kv2.1 channels. Together, these findings demonstrate that the Kv5.1 electrically silent subunit is a component of a substantial fraction of native brain Kv2 channels, and that its incorporation into heteromeric channels can impact diverse aspects of Kv2 channel function.

## Introduction

The Kv2 family of voltage-gated K^+^ channels are highly expressed in brain and play important canonical roles as voltage-activated K^+^-selective channels that regulate neuronal action potentials and membrane excitability (Guan et al., 2007; McKeown et al., 2008; Liu and Bean, 2014; Trimmer, 2015). Indeed, Kv2.1 KO mice are hyperactive, susceptible to seizures, and have enhanced neuronal activity (Speca et al., 2014), and patients with *de novo* mutations in Kv2.1 have epileptic encephalopathy (Allen et al., 2020). In addition, Kv2 channels are highly clustered on neuronal cell bodies and proximal dendrites and have a separate structural role in organizing endoplasmic reticulum (ER) - plasma membrane (PM) junctions (Fox et al., 2015; Kirmiz et al., 2018b). These sites are formed by Kv2 channels in the PM binding to VAP proteins in the underlying ER (Johnson et al., 2018; Kirmiz et al., 2018a) in a phosphorylation dependent manner (Lim et al., 2000; Kirmiz et al., 2018a), making these structures dynamic and responsive to several types of stimuli (Misonou et al., 2004; Misonou et al., 2005). Numerous signaling proteins are recruited to Kv2-containing ER-PM junctions, including L-type Ca^2+^ channels and an array of Ca^2+^- regulated signaling proteins (Vierra et al., 2019), lipid-handling proteins (Kirmiz et al., 2019; Sun et al., 2019), and protein kinase A (PKA) signaling proteins (Vierra et al., 2023). Thus, these microdomains are thought to serve as important hubs for somatodendritic Ca^2+^, lipid, and PKA signaling in brain neurons (Kirmiz et al., 2019; Vierra et al., 2019; Vierra et al., 2021; Vierra et al., 2023).

The Kv2 family consists of only two voltage sensing and pore forming α subunits (Kv2.1 and Kv2.2) (McKeown et al., 2008; Trimmer, 2015). Kv2.1 and Kv2.2 are robustly expressed in multiple neuron types in cortex and hippocampus (Trimmer, 1991; Maletic-Savatic et al., 1995; Bishop et al., 2015), and form both homo- and hetero-tetrameric channels, which have similar functional and structural properties (Kihira et al., 2010). Another potential source of Kv2 channel diversity, however, comes from the ten modulatory, or “electrically silent” K^+^ channel (KvS) subunit genes (Bocksteins et al., 2009; Bocksteins and Snyders, 2012; Bocksteins, 2016). The KvS subunits (Kv5.1, Kv6.1-6.4, Kv8.1-8.2, and Kv9.1-9.3) have sequence similarity to Kv2 subunits, but studies in heterologous cells have shown that KvS subunits do not form functional channels by themselves. Rather, they obligately co-assemble with Kv2 subunits to form heterotetrameric channels which have distinct biophysical properties (Bocksteins and Snyders, 2012). The precise composition of KvS/Kv2 channels remains uncertain, with evidence for dominant Kv2:KvS stoichiometries of either 2:2 (Moller et al., 2020) or 3:1 (Kerschensteiner et al., 2005; Pisupati et al., 2018). Interestingly, KvS mRNAs are expressed in tissue and cell-specific manners that partially overlap with Kv2.1 and Kv2.2 expression (Castellano et al., 1997; Salinas et al., 1997b; Kramer et al., 1998; Bocksteins and Snyders, 2012; Bocksteins, 2016), suggesting that KvS subunits could generate diverse Kv2 channel subtypes, which modulate Kv2 function in a neuron-specific manner. Consistent with this, genetic mutations and gene targeting studies have linked disruptions in the function of KvS-containing channels to epilepsy (Jorge et al., 2011), neuropathic pain sensitivity (Tsantoulas et al., 2018), labor pain (Lee et al., 2020) and retinal cone dystrophy (Wu et al., 2006; Hart et al., 2019; Inamdar et al., 2022), stressing their functional importance in specific cell types. However, there is currently no information available on the contribution of KvS proteins to native Kv2 channels in brain.

In this study, we used mass spectrometry-based proteomics analyses to identify several KvS subunits as components of native Kv2 channels in brain, with Kv5.1 being the most abundant. We find that Kv5.1 is a relatively common subunit of brain Kv2 channels, and that Kv5.1 protein is highly expressed in select layers of cortex. Moreover, Kv2/Kv5.1 heteromeric channels localize at ER-PM junctions on neuronal somata and proximal dendrites. This provides direct evidence that neuron-specific expression of KvS subunits creates diversity in Kv2 channels in brain, with likely impacts on their function.

## Materials and Methods

### Antibodies

All primary antibodies used in this study are listed in Supplementary Table ^1^. Anti-Kv5.1 rabbit polyclonal (pAb) antibodies were generated for this study. In brief, rabbits were immunized with a recombinant fragment corresponding to the C-terminal 76 amino acids (a.a. 419-494) of human Kv5.1 (Uniprot Q9H3M0) produced in E. coli (Genscript, Piscataway, NJ). This C-terminal sequence is unrelated to the C-termini of Kv2.1 and Kv2.2 and is also highly divergent between KvS family members. Rabbit antisera was generated at Pocono Rabbit Farm (Canadensis, PA). Anti-Kv5.1 pAbs were affinity purified from the antisera by strip purification against nitrocellulose membranes onto which 2 mgs of the recombinant Kv5.1 protein had been electrophoretically transferred from an SDS curtain gel (Olmsted, 1981). The anti-Kv5.1C pAb was validated by immunocytochemistry and immunoblotting of HEK cells transfected with recombinant Kv5.1, and its specificity confirmed by a lack of immunolabeling of parallel samples of HEK cells transfected with Kv2.1, Kv2.2 and other KvS family members.

### DNA constructs

Plasmids encoding untagged and GFP-, DsRed- or HA-tagged Kv2.1 and Kv2.2 have been previously described (Lim et al., 2000; Bishop et al., 2018; Kirmiz et al., 2018a). Plasmid encoding human Kv5.1 was generously provided by Dr. Dirk Snyders (Stas et al., 2015), and a plasmid encoding human Kv5.1 with a C-terminal GFP tag was generated by Genscript. The Kv5.1BBS construct was generated at Genscript by inserting a bungarotoxin-binding site (GGWRYYESSLLPYPDGG) at amino acid 215 in the external S1-S2 loop of Kv5.1, as well as an HA tag at the C-terminus. DsRed-Kv2.1BBS was generated at Genscript using an analogous approach with the BBS inserted at amino acid 218 in the S1-S2 loop of rat Kv2.1 (Uniprot P15387). The DsRed-Kv2.1 S586A plasmid has been described previously (Kirmiz et al., 2018a). Plasmids encoding mouse KvS subunits with C-terminal myc-DDK tags were obtained from Origene (Kv6.1 (MR223857); Kv6.4 (MR224440); Kv9.1 (MR218800) and Kv9.2 (MR207651). Plasmid encoding rat Kv8.1 with a C-terminal HA tag was generated by Twist Bioscience. Human Navβ2 plasmid was a kind gift from Dr. Alfred George (Lossin et al., 2002).

### Animals

All procedures using mice and rats were approved by the University of California, Davis Institutional Animal Care and Use Committee and performed in accordance with the NIH Guide for the Care and Use of Laboratory Animals. Animals were maintained under standard light-dark cycles and allowed to feed and drink ad libitum. Adult C57BL/6 J mice 12-16 weeks old of both sexes were used in immunohistochemistry and proteomic experiments. Kv2.1-KO mice (RRID:IMSR_MGI:3806050) have been described previously (Jacobson et al., 2007; Speca et al., 2014), and were generated from breeding of Kv2.1^+/-^ mice that had been backcrossed on the C57/BL6J background (RRID:IMSR_JAX:000664). All experiments with Kv2.1-KO mice used WT littermates as controls. Kv2.2-KO mice were obtained from Drs. Tracey Hermanstyne and Jeanne Nerbonne, and have been described previously (Hermanstyne et al., 2010; Hermanstyne et al., 2013). Kv2.1 and Kv2.2 double-KO (Kv2 dKO) mice (Kv2.1−/−/Kv2.2−/−) were generated by breeding Kv2.1+/− mice with Kv2.2−/− mice.

### Proteomics

Three pairs of WT and Kv2.1 KO mouse littermates were used to prepare DSP-crosslinked mouse brain samples for immunopurifications. Mice were acutely decapitated in the absence of anesthesia and their brains were rapidly removed. Excised brains were homogenized over ice in a Dounce homogenizer containing 5 mL homogenization and crosslinking buffer (in mM): 320 sucrose, 5 NaPO4, pH 7.4, supplemented with 100 NaF, 1 PMSF, protease inhibitors, and 1 DSP (Lomant’s reagent, ThermoFisher Cat# 22585). Following a 1-h incubation on ice, DSP was quenched with 20 mM Tris, pH 7.4 (JT Baker Cat# 4109-01 [Tris base]; and 4103-01 [Tris-HCl]). 2 mL of this homogenate was then added to an equal volume of ice-cold 2× radioimmunoprecipitation assay (RIPA) buffer (final concentrations): 1% (vol/vol) TX-100, 0.5% (wt/vol) deoxycholate, 0.1% (wt/vol) SDS, 150 NaCl, 50 Tris, pH 8.0 and incubated for 30 min at 4 °C on a tube rotator. Insoluble material was then pelleted by centrifugation at 12,000 × g for 10 min at 4 °C. The supernatant was incubated overnight at 4 °C with the anti-Kv2.1 rabbit polyclonal antibody KC (Trimmer, 1991) and then incubated with 100 μL of magnetic protein G beads (ThermoFisher Cat# 10004D) for 1 h at 4 °C on a tube rotator. Beads were then washed 6 times following capture on a magnet in ice-cold 1× RIPA buffer, followed by four washes in 50 mM ammonium bicarbonate (Sigma Cat# A6141). Proteins captured on magnetic beads were digested with 1.5 mg/mL trypsin (Promega Cat# V5111) in 50 mM ammonium bicarbonate overnight at 37 °C. Non-cross-linked mouse brain samples were prepared using similar methodology, except that DSP was omitted from the homogenization buffer. Immunoprecipitations were then performed using anti-Kv2.1 (KC) and anti-Kv5.1 antibodies (Kv5.1C), as described above. Proteomic profiling was performed at the University of California, Davis Proteomics Facility. Tryptic peptide fragments were analyzed by LC-MS/MS on a Thermo Scientific Q Exactive Plus Orbitrap Mass spectrometer as described previously (Kirmiz et al., 2018a; Vierra et al., 2019; Vierra et al., 2023). The proteomics data in this study have been deposited to the ProteomeXchange Consortium via the PRIDE partner repository with the dataset identifier PXD044574 (Vierra et al., 2023).

### Immunoprecipitations and Immunoblotting

For immunoblot analyses of Kv2 and Kv5.1 channels in brain, mouse brains were removed rapidly and homogenized in ice-cold homogenization buffer (described above). Samples of homogenate were extracted by adding an equal volume of 2x RIPA buffer for 15 min, and insoluble material was pelleted by centrifugation at 12,000 x g for 10 min at 4 °C. For IPs, antibodies were added to the supernatant and incubated for 2-3 hr at 4 °C on a rotator or rocking platform, and then magnetic protein G beads (ThermoFisher Cat# 10004D) were added for an additional 1 hr. After washing 3x in 1x RIPA buffer, the IPed proteins were eluted in 2x SDS LB at 70 °C for 5 min. Where appropriate, samples of the lysate were taken before and after each IP reaction to monitor its efficiency. All samples were separated on 8% SDS-PAGE gels and transferred to PVDF membrane. Immunoblots were blocked in LiCor Intercept PBS blocking buffer (LiCor Biosciences) and probed for Kv2.1 (K89/34 mAb or KC pAb), Kv2.2 (N372B/60 mAb or Kv2.2C pAb) and Kv5.1 (Kv5.1C pAb). Grp75/Mortalin (mAb N52A/42) was used as a loading control. Dye-conjugated fluorescent secondary antibodies (LiCor Biosciences) were used to detect bound primary antibodies and imaged using an Odyessy DLx scanner (LiCor Biosciences). A dye-conjugated light chain-specific anti-rabbit secondary antibody (Jackson Labs) was used to detect bound Kv5.1 antibody, to avoid recognition of rabbit antibody heavy chain, whose molecular weight is similar to Kv5.1. Immunoblots were analyzed using Image Studio software (LiCor Biosciences) and statistical analysis was performed using GraphPad Prism. Immunoblot analyses of Kv2 and Kv5.1 channels expressed in HEK cells was performed in a similar manner.

### HEK293T cell culture and transfection

HEK293T cells (ATCC Cat# CRL-3216) were maintained in Dulbecco’s modified Eagle’s medium (Gibco Cat# 11995065) supplemented with 10% Fetal Clone III (HyClone Cat# SH30109.03), 1% penicillin/ streptomycin, and 1x GlutaMAX (ThermoFisher Cat# 35050061) in a humidified incubator at 37 °C and 5% CO2. Cells were transiently transfected using Lipofectamine 3000 (Life Technologies, cat. #L3000008;) following the manufacturer’s protocol, then rescued in fresh growth medium, and used for experiments 36-48 hours after transfection. For immunolabeling, cells were plated and transfected on poly-l-lysine (Sigma Cat# P1524) -coated microscope cover glasses (VWR Cat# 48366-227). For immunoblotting, cells were grown and transfected on 60 mm tissue culture dishes.

### CHO-K1 cell culture and transfection

The CHO-K1 stable cell line expressing a tetracycline-inducible rat Kv2.1 construct (Kv2.1-CHO; (Trapani and Korn, 2003)) was cultured as described previously (Tilley et al., 2014). Cells were transiently transfected using Lipofectamine 3000 for 6 h, rescued in fresh media, and then Kv2.1 expression was induced in Kv2.1-CHO cells with 1 μg/ml minocycline (Enzo Life Sciences, cat. #ALX-380-109-M050;), prepared in 70% ethanol at 2 mg/ml. Voltage clamp recordings were performed 12-24 hours later. During recordings experimenter was blinded to which cells had been transfected with Kv5.1 or Navβ2.

### Electrophysiology

Voltage clamp was achieved with a dPatch amplifier (Sutter Instruments) run by Sutterpatch (Sutter Instruments). Solutions for Kv2.1-CHO cell voltage-clamp recordings: CHO-internal (in mM) 120 K-methylsulfonate, 10 KCl, 10 NaCl, 5 EGTA, 0.5 CaCl2, 10 HEPES, 2.5 MgATP pH adjusted to 7.2 with KOH, 289 mOsm. CHO-external (in mM) 145 NaCl, 5 KCl, 2 CaCl2, 2 MgCl2, 10 HEPES pH adjusted to 7.3 with NaOH, 298 mOsm. Osmolality was measured with a vapor pressure osmometer (Wescor #5520). The calculated liquid junction potential for the internal and external recording solutions was 9.7 mV and not accounted for. For voltage-clamp recordings, Kv2.1-CHO cells were detached in a PBS-EDTA solution (Gibco Cat# 15040-066), spun at 500 g for 2 minutes and then resuspended in 50% cell culture media and 50% CHO-external recording solution. Cells were then added to a recording chamber (Warner Cat# 64–0381) and were rinsed with the CHO-external patching solution after adhering to the bottom of the recording chamber. Transfected Kv2.1-CHO cells were identified by GFP fluorescence and were selected for whole cell voltage clamp. Thin-wall borosilicate glass recording pipettes (Sutter Cat# BF150-110-10) were pulled with blunt tips, coated with silicone elastomer (Sylgard 184, Dow Corning), heat cured, and tip fire-polished to resistances less than 3 MΩ. Series resistance of 2–14 MΩ was estimated from the whole-cell parameters circuit. Series resistance compensation between 13 and 90% was used to constrain voltage error to less than 15 mV, lag was 6 µs. Capacitance and Ohmic leak were subtracted using a P/4 protocol. Output was low-pass filtered at 5 kHz using the amplifier’s built-in Bessel and digitized at 25 kHz. Experiments were performed on Kv2.1-CHO cells with membrane resistance greater than 1 GΩ assessed prior to running voltage clamp protocols while neurons were held at a membrane potential of –80 mV. A stock of 1 mM RY785 (Cayman Cat# 19813) in 10% DMSO which had been stored at -20 °C was kept on ice and diluted in room temperature (20–22°C) CHO-external solution just prior to application. Solutions were flushed over the voltage clamped cell at a rate of approximately 1 mL/min and then flow was stopped for recording. Kv2.1-CHO cells were given voltage steps from -80 mV to 0 mV for 200 ms every 6 seconds during application of RY785 until currents stabilized. When vehicle control was applied to cells, 0 mV steps were given for a similar duration. DMSO concentration in RY785 and vehicle control was 0.01%. Perfusion lines were cleaned with 70% ethanol then milliQ water. Conductance values were determined from tail current levels at 0 mV after 200 ms steps to the indicated voltage. Tail currents were mean current amplitude from 1 to 5 ms into the 0 mV step. Conductance-voltage relations were fit with a Boltzmann function:

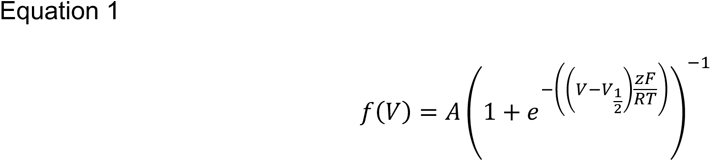

Where V = voltage, A = the amplitude, z = number of elementary charges, F = Faraday’s constant, R = the universal gas constant, and T = temperature = 295 K.

### Immunolabeling of cells

HEK293T cells were fixed for 15 minutes at 4 °C either in 4% formaldehyde prepared fresh from paraformaldehyde in PBS buffer pH 7.4, or in 2% formaldehyde prepared fresh from paraformaldehyde in Na acetate buffer pH 6.0. All subsequent steps were performed at RT. Cells were then washed 3 x 5 minutes in PBS, followed by blocking in blotto-T (Tris-buffered saline [10 mM Tris, 150 mM NaCl, pH 7.4] supplemented with 4% (w/v) non-fat milk powder and 0.1 % (v/v) Triton-X100 [Roche Cat# 10789704001]) for 1 hour. Cells were immunolabeled for 1-2 hours with primary antibodies diluted in blotto-T and subsequently washed 3 x 5 minutes in blotto-T. They were then incubated with mouse IgG subclass- and/or species-specific Alexa-conjugated fluorescent secondary antibodies (Invitrogen) diluted in blotto-T for 45 min and washed 3 x 5 minutes in PBS. Cover glasses were mounted on microscope slides with Prolong Gold mounting medium (ThermoFisher Cat # P36930) according to the manufacturer’s instructions.

For cell surface immunolabeling of HEK293T cells, live cells were incubated for 30 min in 2 ug/ml (250 nM) AF647-alpha-Btx diluted in normal growth medium. Cells were then washed 2x in PBS without Triton X-100, followed by fixation with 2% formaldehyde prepared in PBS for 15 minutes. Cells were then washed 3 x 5 minutes in PBS and processed for labeling of intracellular proteins as described above.

For TIRF imaging, HEK293T cells were plated at low density (30,000-50,000 cells/dish) on 35 mm glass bottom dishes (MatTek #P35G-1.5-14-C) coated with poly-L-lysine (Sigma #P1524) and transfected with DsRed-Kv2.1 BBS and Kv5.1-GFP, GFP-Kv2.1 S586A, or pEGFP-N1. Approximately 48 hours after transfection, cells were incubated for 20 minutes at 37 °C with 1µg/mL α-bungarotoxin-AlexaFluor647 (Invitrogen #B35450) dissolved in HEK293T cell culture medium, followed by a single wash in a modified Krebs-Ringer buffer (KRB) containing (in mM): 146 NaCl, 4.7 KCl, 2.5 CaCl2, 0.6 MgSO4, 1.6 NaHCO3, 0.15 NaH2PO4, 8 glucose, 20 HEPES, pH 7.4. TIRF images of transfected cells in KRB were acquired at 37 °C using the TIRF module of a Marianas SDC Real Time 3D Confocal-TIRF microscope (Intelligent Imaging Innovations) fitted with a 100×, 1.46 NA objective and a Photometrics QuantEM EMCCD camera controlled by Slidebook software (Intelligent Imaging Innovations, 3i). Analysis of the CV of fluorescence intensity was performed using Fiji (NIH). Images for presentation were exported as TIFFs and linearly scaled for min/max intensity and flattened as RGB TIFFs in Photoshop (Adobe).

### Immunolabeling of brain sections

Following administration of sodium pentobarbital (Nembutal, 60 mg/kg) to induce deep anesthesia, animals were transcardially perfused with 4% formaldehyde (freshly prepared from paraformaldehyde) in 0.1 M sodium phosphate buffer pH 7.4, or with 2% formaldehyde (freshly prepared from paraformaldehyde) in 0.05M Na acetate buffer pH 6.0. Sagittal brain sections (30-50 µm thick) were prepared and immunolabeled using free-floating methods as detailed previously (Rhodes et al., 2004; Speca et al., 2014; Bishop et al., 2015; Palacio et al., 2017). Sections were permeabilized and blocked in 0.1 M PB containing 10% goat serum and 0.3% Triton X-100 (vehicle) for 1 hour at RT, then incubated overnight at 4 °C in primary antibodies (Table 2) diluted in vehicle. All subsequent steps were performed at RT. After 4 x 5 min washes in 0.1 M PB, sections were incubated with mouse IgG subclass- and/or species-specific Alexa-conjugated fluorescent secondary antibodies (Invitrogen) and Hoechst 33258 DNA stain diluted in vehicle for 1 hour. After 2 x 5 minute washes in 0.1 M PB followed by a single 5 minute wash in 0.05 M PB, sections were mounted and air dried onto gelatin-coated microscope slides, treated with 0.05% Sudan Black (EM Sciences) in 70% ethanol for 2 minutes (Schnell et al., 1999). Samples were then washed extensively in water and mounted with Prolong Gold (ThermoFisher Cat # P36930).

**Table 1.** Antibody Information.

| **Antigen and Antibody** | **Immunogen** | **Manufacturer information** | **Concentration used** | **Figures** |
| --- | --- | --- | --- | --- |
| **Kv2.1 (KC)** | Synthetic peptide aa 837-853 of rat Kv2.1 | Rabbit pAb, In-house (Trimmer Laboratory), RRID:AB_2315767 | Affinity purified, 1:100 | 1,2 |
| **Kv2.1 (K89/34)** | Synthetic peptide aa 837-853 of rat Kv2.1 | Mouse IgG1 mAb, NeuroMab, RRID:AB_1067225 | Purified, 2 ug/ml | 2 - 7 |
| **Kv2.2 (Kv2.2C)** | Fusion protein aa 717-907 of rat Kv2.2 | Rabbit pAb, in-house (Trimmer Laboratory), RRID:AB_2801484 | Affinity purified, 1:100 | 2,3 |
| **Kv2.2 (N372B/60)** | Fusion protein aa 717-907 of rat Kv2.2 long isoform | Mouse IgG2a mAb, NeuroMab, RRID:AB_2315867 | Purified, 10 µg/mL | 4 - 7 |
| **Kv5.1 (5.1C)** | Fusion protein aa 419-494 of human Kv5.1 | Rabbit pAb, In-house (Trimmer Laboratory) RRID:AB_3076240 | Affinity purified, 1:100 | 1 - 7 |
| **Kv1.2 (1.2C)** | Synthetic peptide aa 463-480 of rat Kv1.2 | Rabbit pAb, In-house (Trimmer Laboratory), RRID:AB_2756300 | Affinity-purified, 1:100 | 2 |
| **Grp75/Mortalin (N52A/42)** | n/a | Mouse IgG1 mAb, NeuroMab, RRID:AB_10674108 | Tissue culture supernatant, 1:10 | 3 |
| **Anti-HA (12CA5)** | Human influenza virus hemagglutinin protein aa 98-106 | Mouse IgG2b mAb, RRID:AB_2532070 | Purified, 5 µg/mL | 1 |
| **Anti-GFP** | GFP Fusion protein | Rabbit pAb Rockland #600-401-215 | Affinity purified, 1.25 ug/ml | 1 |

### Image acquisition and analysis

Widefield fluorescence images were acquired with an AxioCam MRm digital camera installed on a Zeiss AxioImager M2 microscope or with an AxioCam HRm digital camera installed on a Zeiss AxioObserver Z1 microscope with a 63×/1.40 NA Plan-Apochromat oil immersion objective and an ApoTome coupled to Axiovision software version 4.8.2.0 (Zeiss, Oberkochen, Germany). High magnification confocal images of brain sections were acquired using a Zeiss LSM880 confocal laser scanning microscope equipped with an Airyscan detection unit and a Plan-Apochromat 63×/1.40 NA oil immersion DIC M27 objective.

Morphological and colocalization analyses of fluorescent and immunolabeled proteins were performed using Fiji software. For analysis of Kv2.1, Kv2.2 and Kv5.1 colocalization, images were subjected to “rolling ball” background subtraction and an ROI was drawn around the cell body. Pearson’s correlation coefficient (PCC) values were then collected using the Coloc2 plugin. Relative clustering of Kv2 and Kv5.1 was measured by quantifying the coefficient of variation (CV) of the fluorescence intensity (fluorescence s.d./mean s.d.) in each cell, with a greater CV indicative of a higher degree of protein clustering (Cobb et al., 2015). For comparison of WT and Kv2.1 KOs, analysis was performed on images of brain sections taken using the same exposure times. For presentation, images were linearly scaled for min/max intensity in Fiji and saved as RGB TIFFs.

### Statistical analysis

Measurements derived from immunoblots and image analysis were imported into GraphPad Prism for statistical analysis and presentation. Reported values are mean ± s.e.m. We used a one-way ANOVA to compare multiple experimental groups, with post-hoc Tukey’s or Sidak’s multiple comparisons tests to determine which individual means differed. For all tests, P values **<**0.05 were considered to be significantly different, and exact *P*-values are reported in each figure or figure legend. At least two sets of animals or three independent cultures were used for all experiments; the number of samples analyzed is stated in each figure or figure legend. Electrophysiology statistical tests were performed in Igor Pro software version 8 (Wavemetrics, Lake Oswego, OR). Independent replicates are individual cells.

## Results

### KvS subunits are components of Kv2.1 channel complexes in brain

To characterize the composition of Kv2 channels and proteins associated with Kv2-containing ER-PM junctions in mammalian brain we performed mass spectrometry-based proteomics analyses of immunopurified Kv2.1 channel complexes. The Kv2.1 channels were immunoprecipitated (IPed) from mouse brain homogenates that were chemically cross-linked with DSP during preparation (as in (Kirmiz et al., 2018a)), which creates intra- and inter-molecular links between lysine residues in close spatial proximity (12 angstrom) of one another. Parallel IPs were performed from wild type (WT) and Kv2.1 knock-out (KO) adult mouse brain samples to identify only those proteins that were isolated specifically with Kv2.1 channels. Protein components of Kv2.1 channel complexes were identified by mass spectrometry in data-dependent acquisition mode as described previously (Kirmiz et al., 2018a; Vierra et al., 2023). Notably, in addition to Kv2.1, Kv2.2, VAPs (Johnson et al., 2018; Kirmiz et al., 2018a), phosphatidylinositol transfer proteins Nir2/3 (Kirmiz et al., 2019), Ca^2+^ signaling proteins (Vierra et al., 2019) and PKA signaling machinery (Vierra et al., 2023), we identified several members of the electrically silent or KvS subunit family. These are listed in Fig 1a by the total unique mass spectra returned, a method known as spectral counting, and are expressed as a percentage of total Kv2.1 spectra. Notably, we found that the spectral abundance of Kv5.1 was ≈18% of the levels of Kv2.1 and ≈50% of the levels of Kv2.2, indicating that this KvS subunit is a relatively common component of these native brain Kv2 channel complexes. Other KvS subunits were less abundant, with their spectral abundance relative to Kv2.1 ranging from ≈5% for Kv8.1 to 2% for Kv9.1. No KvS subunits were detected in immunoprecipitations (IPs) from Kv2.1 KO brain homogenates. These findings indicate that KvS subunits contribute, to varying degrees, to native Kv2 channel complexes in mouse brain.

**Figure 1.**
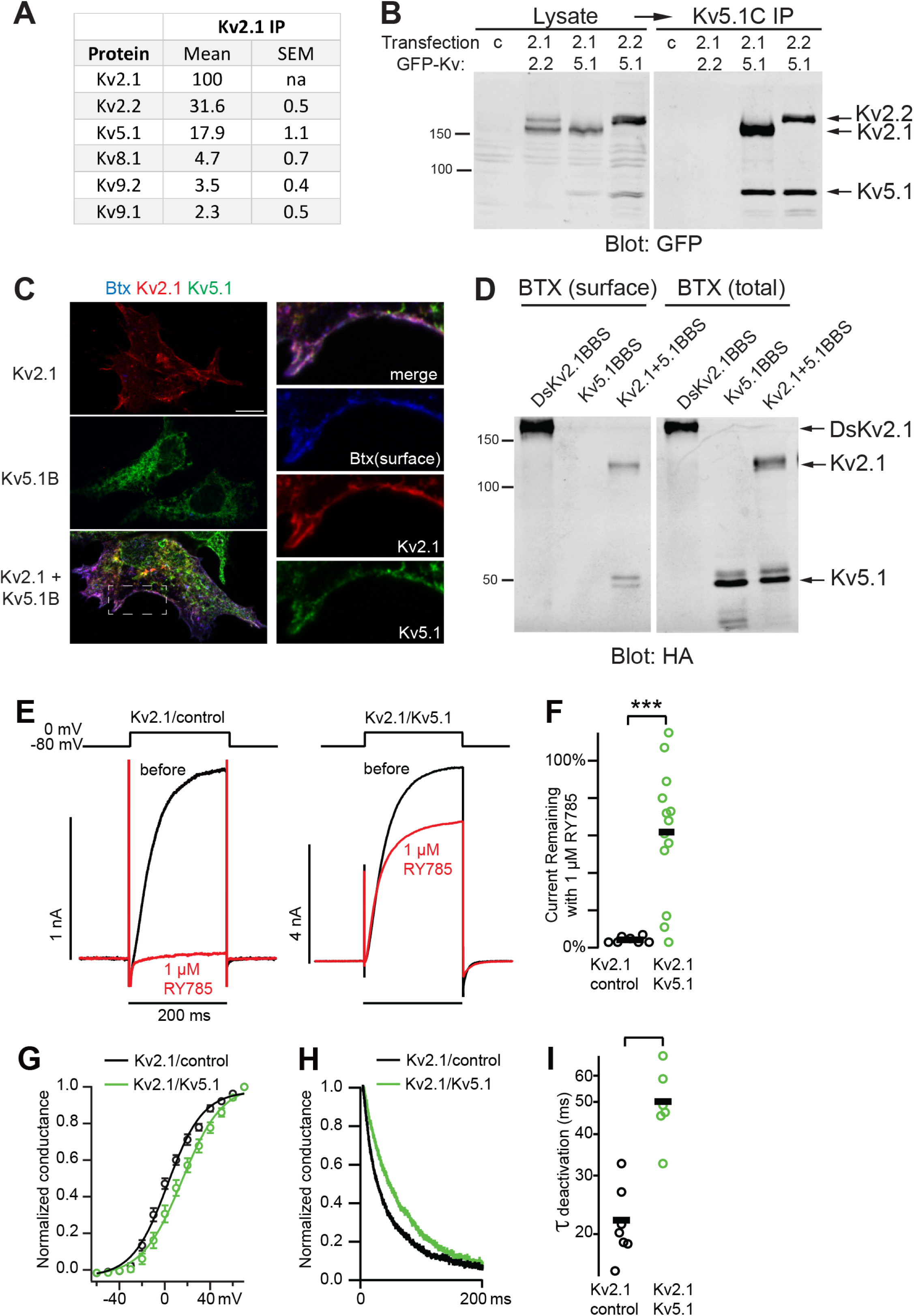
Kv5.1 co-assembles with Kv2 subunits to form heteromeric channels. (A) Mass spectrometry-based proteomics analyses of Kv2.1 complexes immunopurified from crosslinked adult mouse brain samples. The identified proteins are listed by their mean spectral abundance, expressed as a percentage of Kv2.1 spectral counts (mean ± sem, n=3). Several KvS subunits co-purified with Kv2 channels, with Kv5.1 being the most abundant. (B) HEK cells expressing GFP-tagged Kv5.1 and Kv2.1 or Kv2.2, as designated in the lane labels (c = untransfected cells), were solubilized in RIPA buffer and immunoprecipitations (IPs) performed using an affinity-purified Kv5.1 antibody. The IP reactions were size fractionated on SDS gels and immunoblotted for GFP. Kv2.1 and Kv2.2 were both co-IPed with Kv5.1, consistent with their co-assembly into heteromeric channels. Arrows to the right denote positions of the target proteins. Numbers to the left are molecular weights standards in kD. (C) HEK cells expressing Kv2.1 and/or Kv5.1 with an extracellular bungarotoxin binding site (BBS) were cell surface labeled with AF647-Btx (blue), and then permeabilized and immunolabeled with anti-Kv2.1 (red) and Kv5.1 (green) antibodies. Kv5.1BBS was detected on the cell surface with Btx and anti-Kv5.1 antibody only when co-expressed with Kv2.1. Scale bar = 10 μm. (D) HEK cells expressing HA-Kv5.1BBS alone or HA-Kv2.1 + HA-Kv5.1BBS were either labeled live with biotin-Btx (surface) or extracted and then labeled with biotin-Btx (total). The streptavidin precipitation reactions were size fractionated on SDS gels and immunoblotted for the HA epitope tag. Kv5.1 was detected on the cell surface only when co-expressed with Kv2.1. DsRed-HA-Kv2.1BBS is shown as a positive control for surface expression. Arrows to the right denote positions of the IPed proteins. Numbers to the left are molecular weights standards in kD. (E) Exemplar current traces from Kv2.1-CHO cells before (black) and after (green) application of 1 µM RY785. Left panel: transfection control. Right panel: Kv2.1-CHO cells transfected with Kv5.1. (F) Current remaining after application of 1 μM RY785 as in panel A. Black bars represent mean. Each point is current from one cell at the end of a 200 ms 0 mV voltage step. T test *** p<0.001. (G) Voltage dependence of activation normalized to maximum conductance of initial tail currents at 0 mV. Mean ± SEM. Kv2.1/control (black) n = 6 cells, Kv2.1/Kv5.1 (green) n = 7 cells. Lines are Boltzmann fits (Equation 1) Kv2.1/control: V1/2 = 2.6 ± 1 mV z = 1.6 ± 0.1 e0, Kv5.1/Kv2.1: V1/2 = 15 ± 2 mV z = 1.4 ± 0.1 e0. (H) Exemplar traces of channel deactivation at -40 mV after a 50 ms step to +20 mV. Traces are normalized to max current during -40 mV step. (I) The faster time constant of double exponential fits to channel deactivation. T test p = 0.001.

### Kv5.1 and Kv2 subunits form functional channels in heterologous cells

The simplest interpretation of these proteomics data is that KvS subunits co-assemble with Kv2 subunits to form heteromeric channels, although the possibility of indirect cross-linking of independent KvS subunits to Kv2.1 channels cannot be excluded. To better define the role of KvS subunits in brain we next investigated the composition and localization of putative Kv2/KvS channels focusing on Kv5.1 as an exemplar due to its abundance in our proteomics analyses. As a first step, we tested whether Kv5.1 forms heteromeric channels with Kv2.1 and/or Kv2.2 subunits in heterologous cells. For these experiments we utilized a rabbit polyclonal antibody (Kv5.1C) that we generated against the Kv5.1 cytoplasmic C-terminal tail, whose sequence is highly divergent between different KvS subunits. This antibody was validated by ELISA and immunofluorescent labeling, immunoblotting and immunoprecipitation from Kv5.1-transfected cells. Moreover, we confirmed that the antibody did not cross-react to Kv2.1, Kv2.2 (Fig 1) or to other KvS subunits (Suppl. Fig 1). HEK293T (HEK) cells expressing Kv5.1 together with Kv2.1 or Kv2.2 were solubilized in RIPA buffer and IPs performed using the Kv5.1C antibody. Notably, we found that both Kv2.1 and Kv2.2 co-IPed together with Kv5.1 from transfected cell lysates (Fig 1b).

To test whether Kv5.1 co-assembles with Kv2 subunits to form heteromeric channels expressed on the cell surface, we utilized a Kv5.1 construct with a bungarotoxin binding site (BBS) inserted into the extracellular S1-S2 loop. The BBS binds alpha-bungarotoxin (Btx) with high affinity (Sekine-Aizawa and Huganir, 2004) and live cell labeling with Btx thereby allows for specific detection of cell surface-expressed Kv5.1. Indeed, in HEK cells expressing Kv2.1BBS as a positive control, surface Kv2.1 channels were robustly detected with AF647-Btx. In contrast, in HEK cells expressing Kv5.1BBS alone, Kv5.1 was undetectable after surface labeling with AF647-Btx, and immunolabeling with Kv5.1 antibody showed that it was retained intracellularly in an ER-like pattern. When Kv5.1BBS was co-expressed with Kv2.1, however, Kv5.1 was readily detected on the cell surface with AF647-Btx, where it co-localized precisely with Kv2.1 (Fig 1c). Similarly, following live cell labeling with biotin-Btx, Kv5.1BBS was isolated with streptavidin beads from cell lysates only when it was co-expressed with Kv2.1 (Fig 1d). Together, these findings provide strong evidence that Kv5.1 co-assembles with Kv2.1 and Kv2.2 subunits to form heteromeric channels expressed on the cell surface of heterologous cells.

To test whether heteromeric Kv2.1/Kv5.1 channels are functional, we performed electrophysiological recordings. The voltage-gated K^+^ currents from CHO cells stably transfected with Kv2.1 are blocked by the drug RY785 (Marquis and Sack, 2022). We found that transfection of these cells with Kv5.1 resulted in RY785-resistant voltage-gated currents not observed in transfection controls (Fig 1e,f). The RY785-resistant currents of Kv5.1-transfected cells have a conductance-voltage activation relation shifted to more positive voltages (Fig 1g), and slower deactivation (Fig 1h,i), similar to conductances reported after co-injection of Kv2.1 and Kv5.1 into *Xenopus* oocytes (Kramer et al., 1998). We conclude that channels containing these Kv5.1 subunits can be functional at the surface of mammalian cells and have biophysical and pharmacological properties distinct from Kv2.1 homomers.

### Kv5.1 is a common component of native Kv2 channels in brain

Next, we investigated whether Kv5.1 is a component of native Kv2 channels in brain. Non-cross-linked rat brain membranes or mouse whole brain homogenates were solubilized with RIPA buffer and Kv2/KvS channels were IPed with specific antibodies and their composite subunits analyzed by immunoblotting. In IPs performed using Kv5.1-specific antibody (Kv5.1C) we readily detected both Kv2.1 and Kv2.2 subunits by immunoblotting (Fig 2a). Neither of these subunits were detected in IPs using a control antibody against the axonal Kv1.2 channel. Similarly, in reverse IPs using either Kv2 subunit antibody, we found that Kv5.1 was co-immunoprecipitated with Kv2.1 and to a lesser extent with Kv2.2 subunits (Fig 2b). In contrast, Kv5.1 was not detected in control IPs using anti-Kv1.2 antibodies. As all IPs were performed under stringent detergent extraction conditions (*i.e.,* in RIPA buffer), these findings provide compelling evidence that Kv5.1 is an intrinsic component (*i.e.*, subunit) of native Kv2 channels.

**Figure 2.**
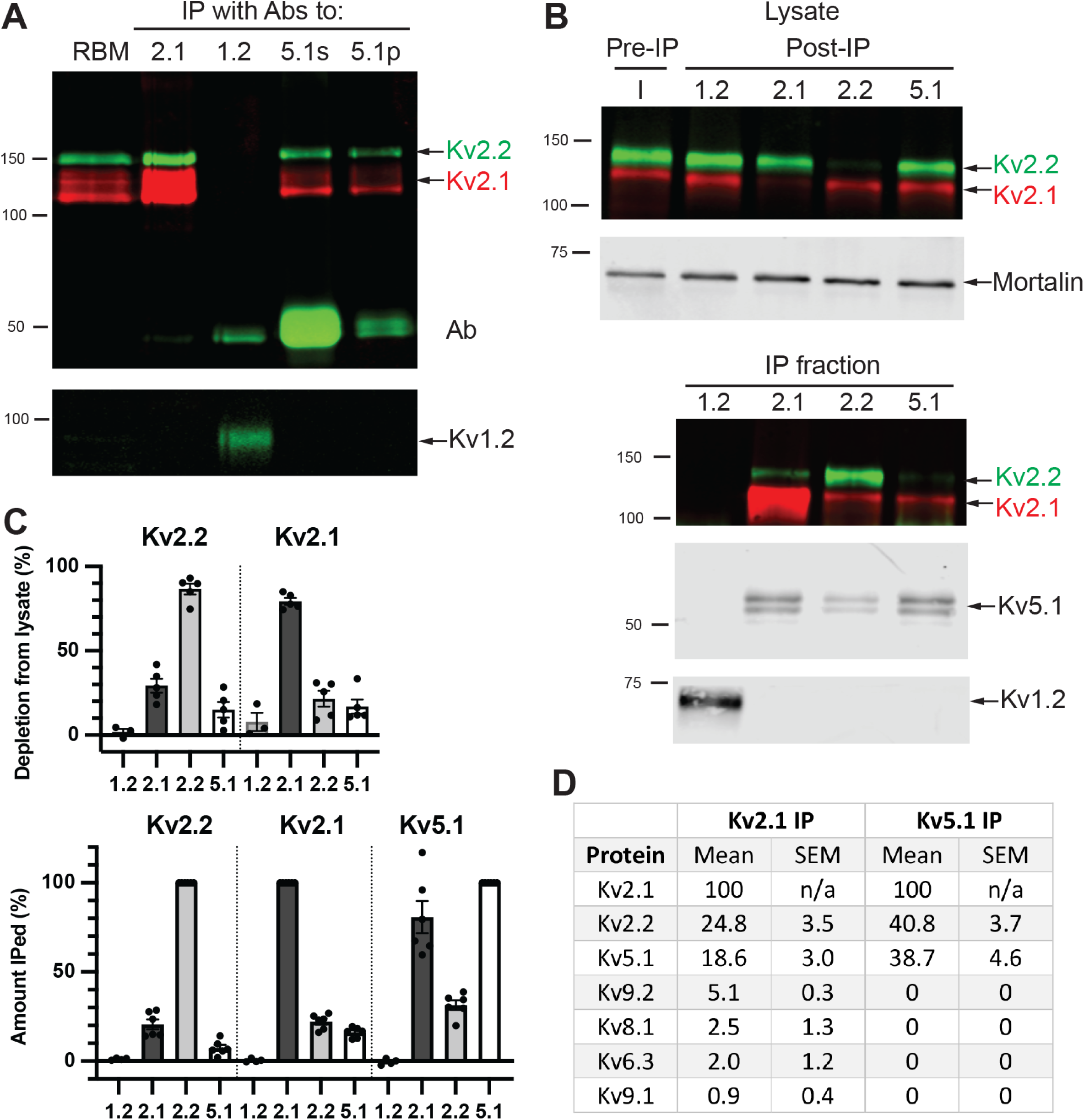
Kv5.1 is a common component of native Kv2 channels in brain. (A) Rat brain membranes (RBM) were solubilized with RIPA buffer and IPs performed using polyclonal antibodies to Kv2.1, Kv1.2, or Kv5.1 (s=serum; p=affinity-purified). The RBM starting material and the reaction products from IPs performed with the various antibodies as designated in the lane labels were size fractionated on SDS gels and immunoblotted for Kv2.2 (green), and Kv2.1 (red). The panel below shows a section of the blot that was probed for Kv1.2. Both Kv2.1 and Kv2.2 were robustly co-IPed together with Kv5.1, but not with Kv1.2. Labels to the right denote positions of the target proteins. Numbers to the left are molecular weights standards in kD. (B) Complementary IPs were performed from mouse brain lysates using Kv2.1 or Kv2.2 subunit antibodies. The input lysate (I), the post-IP depleted lysates (top panels) and the IP fractions (bottom panel) performed with the various antibodies as designated in the lane labels were size fractionated on SDS gels. Immunoblotting of pre- and post-IP lysates (top panel) for Kv2.2 (green), and Kv2.1 (red) confirmed that both IPs were highly efficient. The panel below shows a section of the blot that was probed for GRP75/Mortalin as a loading control. Immunoblotting of the IP fractions (bottom panel) for Kv5.1 shows that it co-IPed together with Kv2.1 and to a lesser extent with Kv2.2, but not with Kv1.2. Labels to the right denote positions of the target proteins. Numbers to the left are molecular weights standards in kD. (C) Quantification of pre- and post-IP brain lysates showed that 87% of Kv2.2 and 79% of Kv2.1 protein were depleted from brain lysates with Kv2.2 and Kv2.1 IPs, respectively (top graph; n=3-5). Quantification of Kv2.2, Kv2.1 and Kv5.1 subunits in each IP, normalized to the amount IPed by each subunit-specific antibody (bottom graph; n=4-6). Approx. 21-22% of Kv2.1 and 2.2 were co-IPed with the other Kv2 subunit, and 16% of Kv2.1 and 7% of Kv2.2 were co-IPed together with Kv5.1. Twice as much Kv5.1 was co-IPed together with Kv2.1 as compared to Kv2.2. (D) Mass spectrometry-based proteomics analyses of Kv2 channels immunopurified from non-cross-linked brain lysates using Kv2.1 and Kv5.1 specific antibodies. Mean spectral abundance is expressed as a percentage of Kv2.1 spectral counts (mean ± sem, n=3). Several KvS subunits were detected in Kv2.1 IPs, with Kv5.1 being the most abundant (18.6% of Kv2.1 levels). In contrast, no other KvS subunits were detected in Kv5.1 IPs.

To estimate the abundance of Kv2/Kv5.1 channels we compared the relative amounts of Kv2.1, Kv2.2 and Kv5.1 subunits isolated in each IP reaction. Immunoblotting of the pre- and post-IP lysates showed that both the Kv2.2 and Kv2.1 IPs are highly efficient, depleting 87% of Kv2.2 and 79% of Kv2.1 from the lysate, respectively (Fig 2c). Consequently, we compared the amount of Kv2.2, Kv2.1 and Kv5.1 subunits in each co-IP to that in each subunit-specific IP. Approximately 21% as much Kv2.2 was isolated in Kv2.1 IPs as compared to Kv2.2 IPs, and 22% as much Kv2.1 was isolated in Kv2.2 IPs as compared to Kv2.1 IPs (Fig 2c). This suggests that over 20% of Kv2.1 and Kv2.2 subunits in brain occur in Kv2.1/Kv2.2 heteromeric channels. By comparison, we found that 7.4% of Kv2.2 and 16% of Kv2.1 were isolated in Kv5.1 IPs, and conversely, 31% of Kv5.1 was isolated in Kv2.2 IPs and 81% in Kv2.1 IPs. Thus, Kv2.1/Kv5.1 heteromeric channels predominate over Kv2.2/Kv5.1 channels in brain.

In a complementary approach we also analyzed Kv2/KvS channel composition using mass spectrometry-based proteomics. For this, IPs were performed from non-cross-linked mouse brain lysates using Kv2.1 or Kv5.1 antibodies and stringent detergent extraction and wash conditions. The identity and spectral abundance of subunits in the Kv2.1- and Kv5.1-containing channels is shown in Fig 2d, expressed as a percentage of total Kv2.1 spectra. In Kv2.1 IPs, Kv2.2 and Kv5.1 subunit spectra were detected at 25% and 19% of Kv2.1 levels, respectively. Consistent with our experiments on crosslinked brain samples, several other KvS subunits (Kv9.2, Kv6.3, Kv8.1 and Kv9.1) were also identified, although with significantly lower spectral counts. These findings suggest that a substantial subpopulation of native Kv2 channels in brain are heteromeric, containing both Kv2.1 and Kv2.2 subunits, and/or Kv2 and KvS subunits. Most notably, the spectral abundance of KvS subunits combined is approximately 30% of that of Kv2.1, indicating that KvS subunits are relatively common components of brain Kv2 channels, and create considerable diversity in this class of Kv channel.

In analogous mass spectrometry analysis of Kv5.1 subunit-containing channels isolated in Kv5.1 IPs, Kv2.1 subunit spectral abundance was over twice that of Kv2.2, supporting our findings from IP and IB experiments (Fig 2d). Moreover, we found that the ratio of Kv5.1 to Kv2.1 plus Kv2.2 subunit spectra (39/(100+41) was ≈1:3.5, although this is likely an underestimate of Kv5.1’s contribution due to its smaller molecular mass compared to Kv2.1 and Kv2.2. Consequently, it is not possible to define the Kv2/KvS channel stoichiometry solely from these data (see Discussion). Interestingly, we did not detect any other KvS subunits in Kv5.1 IPs, suggesting that native brain Kv2/Kv5.1 heterotetrameric channels do not commonly contain multiple KvS subunit types.

### Kv5.1 protein expression is significantly reduced in Kv2 KOs

As Kv5.1 forms heteromeric channels with both Kv2.1 and Kv2.2 subunits in brain, we tested how Kv5.1 expression is impacted in mice with a genetic deletion of one or both Kv2 subunits. We previously found that protein abundance of the Kv2 channel auxiliary subunit AMIGO-1 is significantly depressed in brains from Kv2 KO mice, suggesting that expression of AMIGO-1 in brain depends on its assembly with Kv2 channels [Bishop et al., 2018]. Similar to AMIGO-1, we found by immunoblotting of brain lysates and Kv5.1 IPs that Kv5.1 protein levels are reduced by 70% in Kv2.1 KO mouse brains, and by 95% in Kv2.1/2.2 DKO mouse brains as compared to brains of wild-type mice (Fig 3). The magnitude of these decreases in Kv5.1 expression approximate its relative association with Kv2.1 and Kv2.2 in wild-type brain (Fig 2) and provide further evidence that Kv2.1/Kv5.1 hetero-tetrameric channels predominate over Kv2.2/5.1 channels. Moreover, we found no indication of any compensatory increase in Kv5.1 association with Kv2.2 in Kv2.1 KO brains. Thus, Kv5.1 protein expression depends on Kv2.1 and Kv2.2 subunits, and most likely their co-assembly into heteromeric channels.

**Figure 3.**
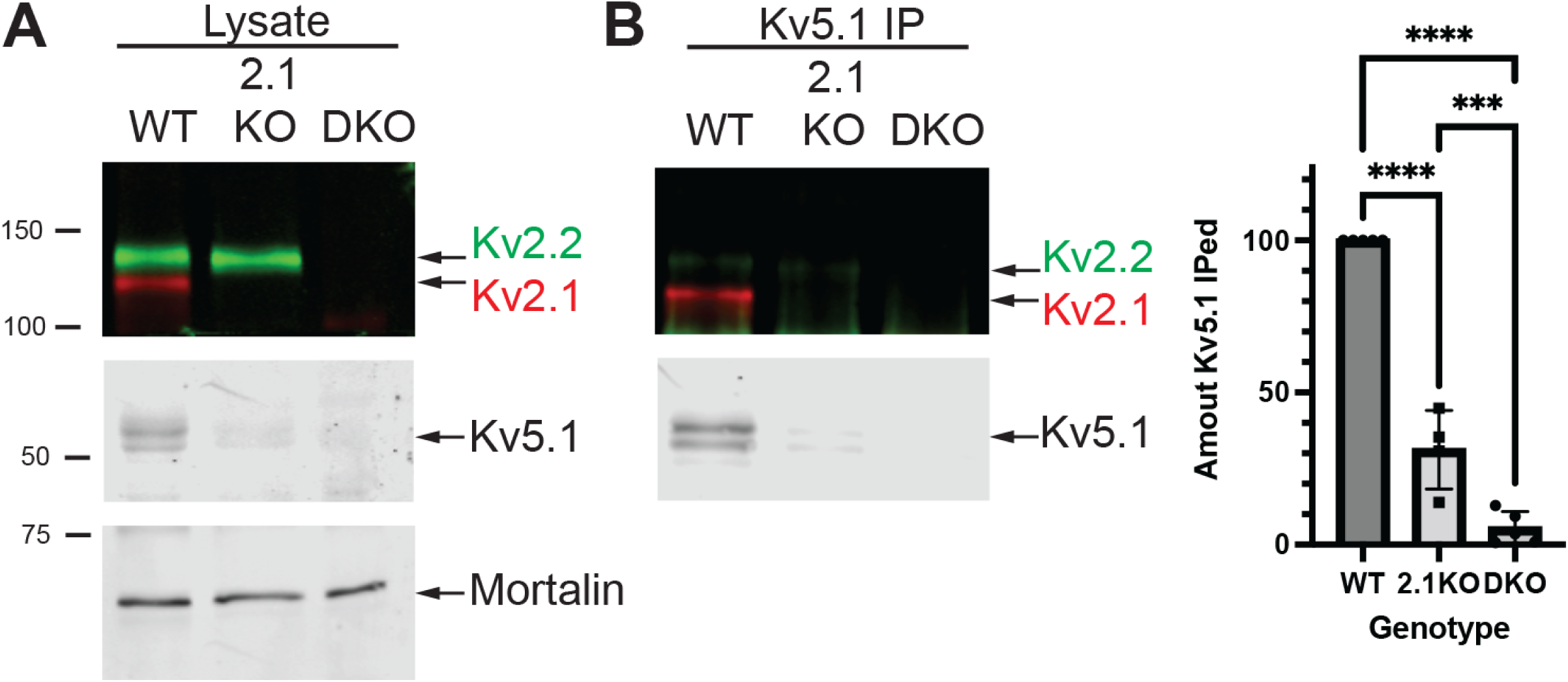
Kv5.1 protein is significantly reduced in Kv2 KO brain. (A) Immunoblots of brain lysates from wild-type (WT), Kv2.1KO and Kv2.1/2.2DKO mice show that Kv5.1 protein levels are severely reduced in Kv2 KO brain. Labels to the right denote positions of the target proteins. Numbers to the left are molecular weights standards in kD. (B) Immunoblots of Kv5.1 IPs from brain lysates of wild-type (WT), Kv2.1KO and Kv2.1/2.2DKO mice show that Kv5.1 protein levels are severely reduced in Kv2 KO brain. Graph shows quantitation of IP reaction products. Compared to WT mouse brain, the amount of Kv5.1 IPed is decreased by 70% in Kv2.1KO and 95% in Kv2.1/2.2DKO brain samples (one way ANOVA and Tukey’s multiple comparisons test, n=4-5, **** p<0.0001). Thus, Kv5.1 expression is dependent on Kv2 subunits and primarily on Kv2.1.

### Kv5.1 protein is highly expressed in rodent cortex

Gene expression studies have shown the KvS family subunit mRNAs are differentially expressed in rodent brain, but their protein expression and localization has remained unknown. Consequently, we defined the regional and cellular patterns of Kv5.1 protein expression in brain using multiplex IF labeling. Notably, we found that effective immunofluorescent labeling of brain sections using the Kv5.1C antibody required specialized fixation conditions or antigen retrieval steps after conventional fixation. Optimal Kv5.1 immunolabeling was obtained in sections from animals that were perfusion-fixed with 2% formaldehyde in pH 6 buffer (Berod et al., 1981). This labeling was specific to the affinity-purified Kv5.1 antibody and not observed using pre-immune serum. Moreover, as shown above (Fig 1), Kv5.1 antibody selectively labeled recombinant Kv5.1 expressed in fixed and permeabilized heterologous cells and did not cross-react to Kv2, or other KvS subunits (data not shown). Importantly, we found substantial regional differences in Kv5.1 protein immunolabeling in brain sections, with high levels of expression observed in cortex and olfactory bulb. Kv5.1 immunolabeling in other regions is largely undetectable with the notable exception of moderate Kv5.1 immunolabeling in the CA3 region of hippocampus. These findings are broadly consistent with in situ hybridization and transcriptomic analysis of Kv5.1 mRNA expression in mouse brain (Allen Brain Atlas). Based on these findings, we focused our detailed analysis of Kv5.1 protein expression and localization in the neocortex.

In the mouse neocortex, multiplex IF labeling of sagittal and coronal brain sections revealed that Kv5.1 immunolabelling varies substantially across different cortical layers (Fig 4). Kv5.1 labeling was most prominent in Layers 2/3, where robust labeling was detected in a substantial subset of neurons that also immunolabeled for Kv2.1 and/or Kv2.2 (Fig 4a,b). Most of these Kv2/Kv5.1 positive neurons appeared to be pyramidal neurons, based on morphological criteria.

**Figure 4.**
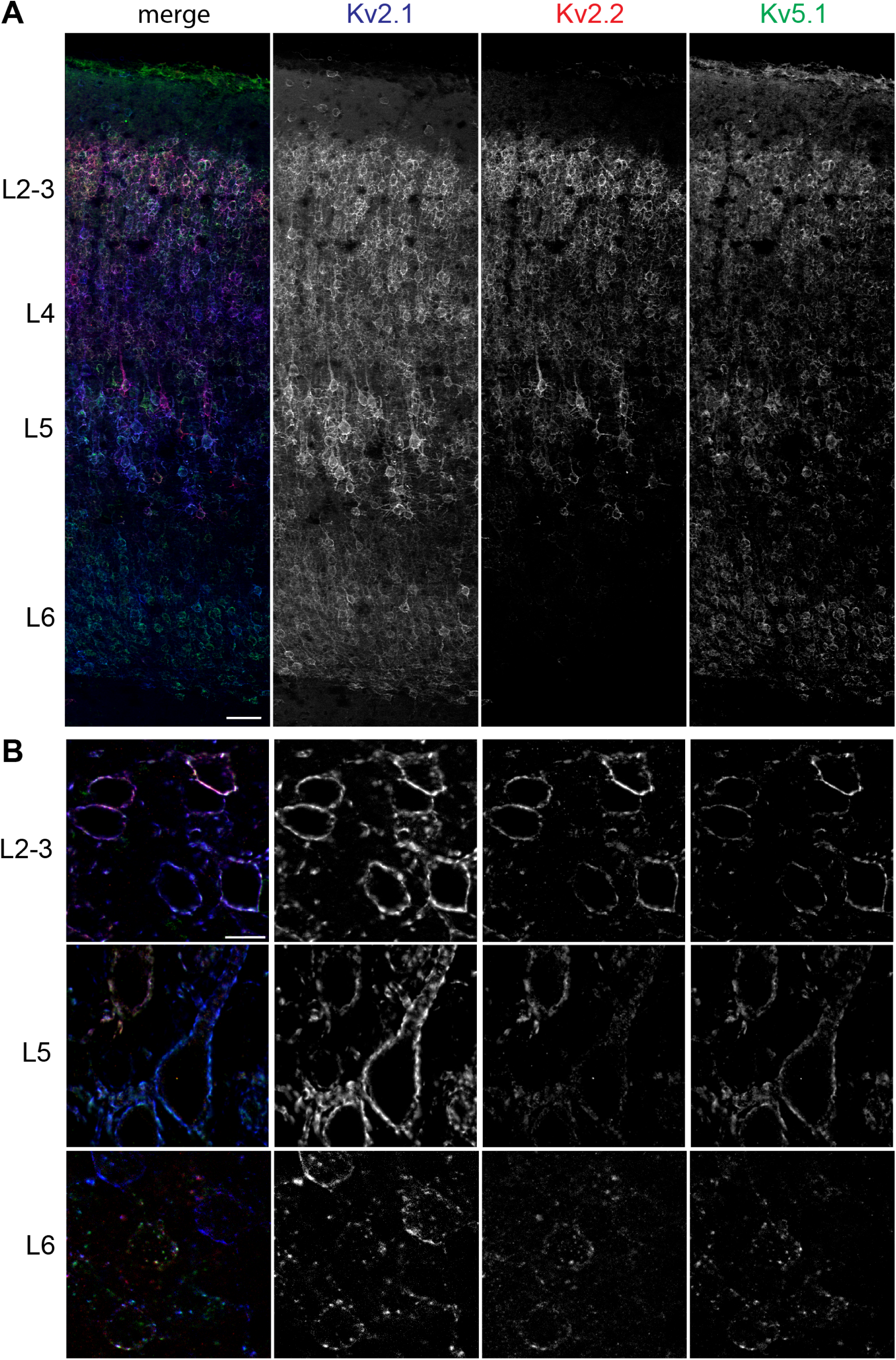
Kv5.1 protein is highly expressed in cortex. (A) Multiplex immunofluorescent labeling of a sagittal section of mouse somatosensory cortex. Kv5.1 (green) immunolabelling varies across cortical layers and is most prominent in layer 2/3, where it is detected in a significant subpopulation of neurons that express Kv2.1 (blue) and/or Kv2.2 (red). Kv5.1 immunolabeling is detected in smaller subpopulations of Kv2 positive neurons in deeper cortical layers. Scale bar = 50 µm. (B) Higher magnification images showing that Kv5.1 immunolabeling is restricted to neurons expressing Kv2.1 and/or Kv2.2, depending on the cortical layer. Scale bar = 10 µm.

By comparison, Kv5.1 immunolabelling was less prominent in layers 4-6 and was detected in a smaller subset of neurons that labeled for Kv2 subunits. Thus, we find that Kv5.1 is differentially expressed across the cortical layers, and that it is restricted to varying subpopulations of Kv2 positive neurons in each layer. Most notably, Kv5.1 is particularly enriched in L2/3, where it is expressed in a large subset of neurons.

### Kv5.1 is co-localized with Kv2 subunits in a subset of cortical neurons

A more detailed analysis in L2/3 of cortex revealed several important features regarding Kv5.1 protein expression. First, we found that Kv5.1 immunolabelling is extensively colocalized at the cellular level with both Kv2.1 and Kv2.2, and that like Kv2 subunits it is prominently expressed on the neuronal somata and proximal dendrites (Fig 5a). As Kv5.1 channels do not contain the VAP-interacting PRC motif required for somatodendritic targeting and clustering (Lim et al., 2000), this immunolabeling pattern is consistent with Kv5.1 incorporation into heteromeric channels containing one or more obligate PRC-domain containing Kv2 α subunits. This is reflected in similar Pearson Correlation Coefficient (PCC) values for Kv2.1 vs Kv5.1 and Kv2.2 vs Kv5.1 immunolabeling, which are only slightly lower than the PCC values for Kv2.1 vs Kv2.2 immunolabeling (Fig 5b). Second, although Kv5.1 immunolabeling colocalizes with that of Kv2.1 and/or Kv2.2, we find that the ratios of Kv2.1, Kv2.2 and Kv5.1 immunolabeling vary between neurons even in the same cortical layer. This is shown qualitatively by the differing hues of neighboring L2/3 neurons immunolabeled for Kv2.1, Kv2.2 and Kv5.1 (Fig 5a), and by the fact that the ratios of total IF labeling intensity between Kv2.1, Kv2.2 and Kv5.1 vary between individual neurons (Fig 5c). This indicates that Kv2.1, Kv2.2 and Kv5.1 protein expression vary between L2/3 neurons, and that Kv2 channel composition can differ considerably even in neighboring neurons. Third, we find that Kv5.1 is co-clustered together with Kv2 channels on neuronal soma and proximal dendrites, at sites known to correspond to ER-PM junctions. This identifies Kv5.1 as a novel component of Kv2-containing ER-PM junctions in many L2/3 cortical neurons.

**Figure 5.**
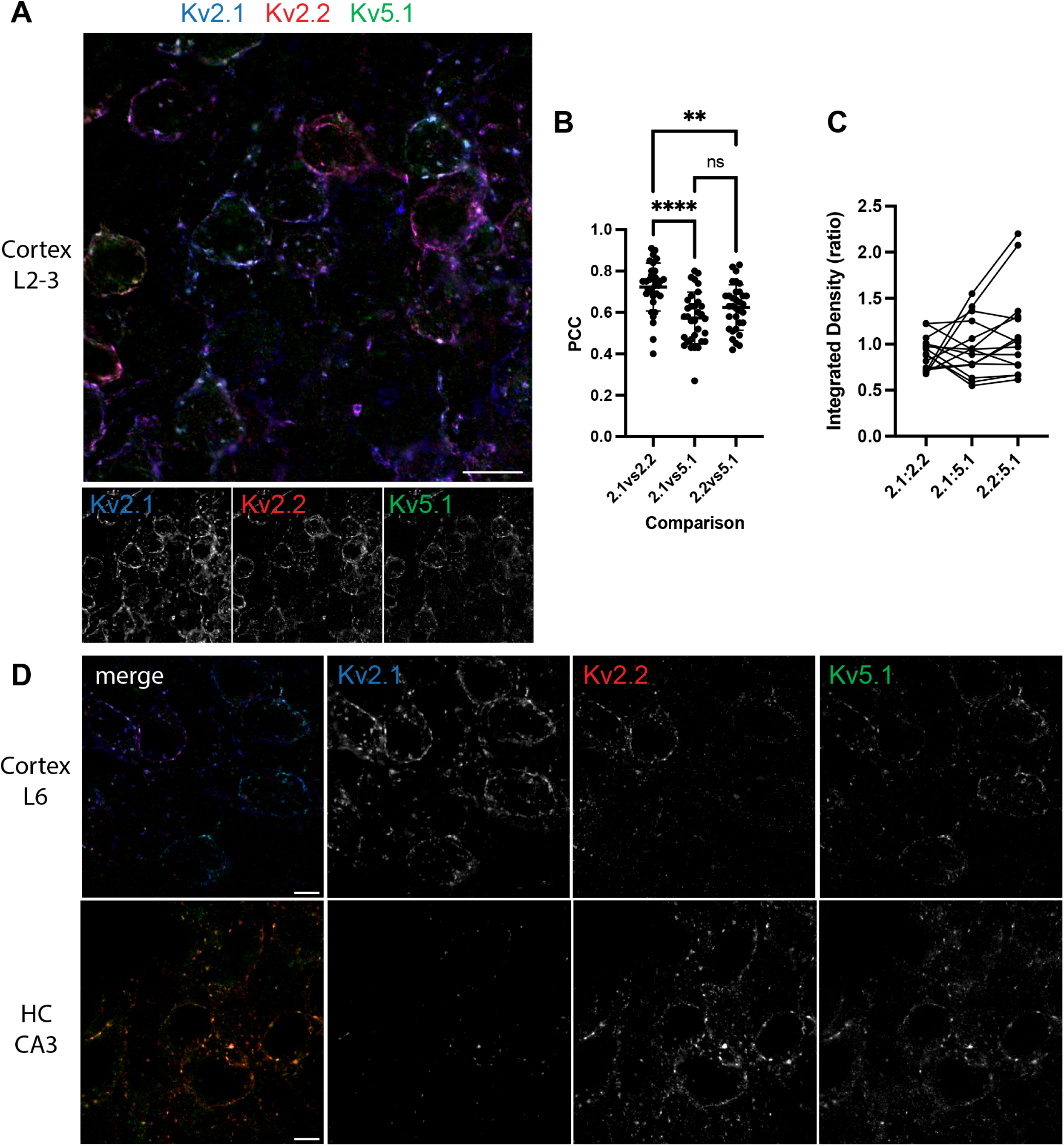
Kv5.1 co-localizes with Kv2 subunits on neuronal somata and proximal dendrites. (A) Confocal image of multiplex immunofluorescent labeling of L2/3 neurons shows that Kv5.1 (green) colocalizes extensively with both Kv2.1 (blue) and Kv2.2 (red) on somata and proximal dendrites. However, the ratios of Kv2.1, Kv2.2 and Kv5.1 immunolabeling vary widely between neighboring neurons (note varying hues of immunolabeling). Scale bar = 10 µm. (B) In L2/3 neurons, similar PCC values demonstrate that Kv5.1 colocalizes equally with Kv2.1 and Kv2.2 (one way ANOVA and Tukey’s multiple comparisons test, **** p<0.0001, ** p=0.0026, n=33 neurons, 3 WT brains). (C) In L2/3 neurons, ratios of Kv2.1, 2.2 and 5.1 immunolabeling intensity vary between neurons (depicted by each line; n=15 neurons). (D) In L6 cortical neurons, Kv5.1 primarily colocalizes with Kv2.1, as Kv2.2 expression is minimal in this layer. In contrast, in CA3 hippocampal neurons, Kv5.1 primarily colocalizes with Kv2.2, as Kv2.1 expression is low in these neurons. Scale bar = 5 µm.

The results obtained for Kv5.1 expression in L2/3 neurons also broadly applies to neurons in other cortical layers. In L4-6, Kv5.1 expression is restricted to a smaller subset of neurons that express Kv2 subunits, and Kv5.1 colocalizes with both Kv2.1 and/or Kv2.2 on their soma and proximal dendrites (Fig 5d). As observed in L2/3 neurons we found considerable heterogeneity in the relative expression levels of Kv2.1, Kv2.2 and Kv5.1 in different neurons. In L6 neurons, however, Kv5.1 colocalized and co-clustered predominantly with Kv2.1, due to the absence of Kv2.2 expression in most neurons in this layer (Fig 5d). Conversely, in CA3 neurons in hippocampus, Kv5.1 co-clustered predominantly with Kv2.2, due to the minimal Kv2.1 expression in those neurons (Fig 5d). These findings provide further evidence that Kv5.1 contributes to distinct Kv2.1 and Kv2.2 channel subpopulations in mammalian brain neurons.

### Kv5.1 immunolabeling is significantly reduced in Kv2.1 KO brain sections

Our biochemical analysis of native Kv2/Kv5.1 channels indicates that Kv5.1 is predominantly associated with Kv2.1 subunits, and that the expression of Kv5.1 is reduced in Kv2.1 KO and Kv2 dKO brain samples, predicts that Kv5.1 immunolabeling should be diminished in Kv2.1 KO brain sections. This is indeed the case, as we found that Kv5.1 immunofluorescent labeling was significantly reduced in cortical neurons in Kv2.1 KO mice, as compared to wild type controls. In L2/3 neurons, both plasma membrane localization and clustering of Kv5.1 were greatly reduced or often undetectable in Kv2.1 KO brains, and residual Kv5.1 immunolabeling was typically co-colocalized with that of Kv2.2 (Fig 6a). Indeed, intensity profile plots show that Kv5.1 still co-clustered with Kv2.2 in some Kv2.1 KO neurons, although the intensity of Kv5.1 immunolabeling was reduced as compared to that in wild-type samples (Fig 6a). Consistent with this, we found that PCC values for Kv2.2 vs Kv5.1 was significantly decreased in in Kv2.1 KO neurons as compared to wild-type neurons (Fig 6b). These findings are consistent with our biochemical analysis (Fig 3) and support that Kv5.1 protein expression depends predominantly on co-assembly with Kv2.1 subunits into heteromeric channels.

**Fig 6.**
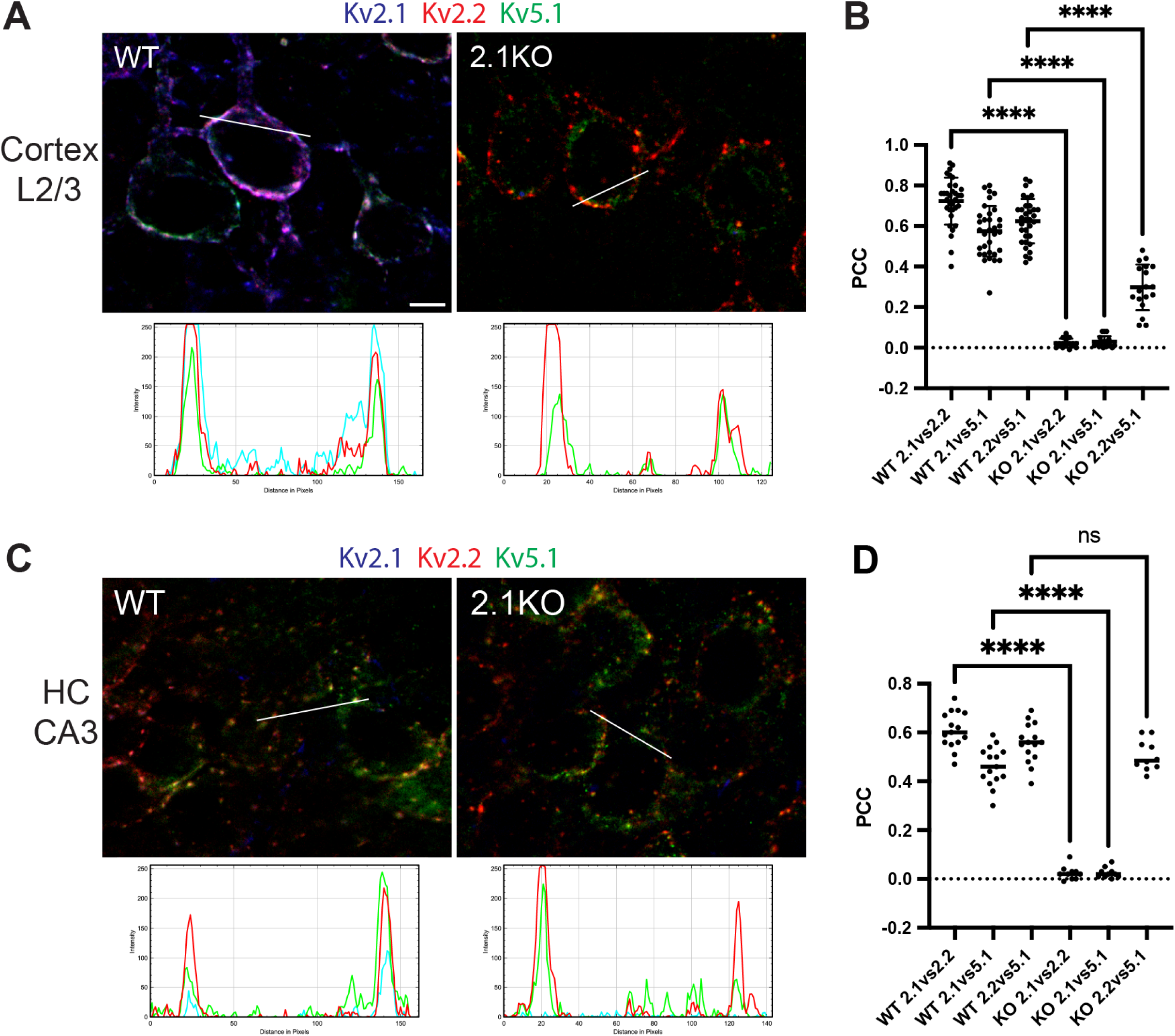
Kv5.1 surface expression is significantly reduced in Kv2.1 KO brain. (A) Immunolabeling for Kv5.1 (green), Kv2.1 (blue) and Kv2.2 (red) in WT (left) and Kv2.1 KO (right) mouse brain sections. Immunolabelling shows that the intensity of Kv5.1 immunolabeling associated with the somatic PM is undetectable or severely reduced in L2/3 neurons in Kv2.1KO brain as compared to wild type (WT). Reduced levels of Kv5.1 remain co-clustered with Kv2.2 in some Kv2.1 KO neurons (as shown in the intensity profile plot). Scale bar = 5 µm. (B) PCC values for Kv2.2 vs. Kv5.1 immunolabelling In L2/3 neurons are significantly reduced in Kv2.1KO brain as compared to WT (one way ANOVA and Sidak’s multiple comparisons test, **** p<0.0001, n=2 pairs of WT and Kv2.1KO brains). (C) Immunolabeling of hippocampal CA3 neurons shows that Kv5.1 is primarily co-clustered with Kv2.2, and that their clustering is preserved in Kv2.1 KO brain. (D) PCCs for Kv2.2 vs. Kv5.1 immunolabelling in CA3 neurons are similar in WT and Kv2.1KO brain (one way ANOVA and Sidak’s multiple comparisons test, **** p<0.0001, n=2).

In one notable exception, we observed grossly normal Kv5.1 expression in the CA3 region of hippocampus in Kv2.1 KO brains (Fig 6c). In wild-type brains, pyramidal neurons in CA3 express significantly higher levels of Kv2.2 compared to Kv2.1 subunits, and correspondingly Kv5.1 exhibits a higher degree of co-localization with Kv2.2 as compared to Kv2.1 (PCC, Fig 6d). In Kv2.1 KO brains, we found that Kv5.1 immunolabeling in CA3 neurons was similar to that in wild-type brains, with Kv2.2 and Kv5.1 remaining co-clustered on neuronal soma and proximal dendrites (Fig 6c). The degree of Kv2.2 and Kv5.1 colocalization in CA3 neurons was also indistinguishable between wild-type and Kv2.1 KO brains (Fig 6d). Thus Kv5.1 expression and localization are selectively preserved in CA3 neurons of Kv2.1 KO mice, likely due to its predominant association with Kv2.2 rather than Kv2.1 subunits in those neurons.

### Kv5.1 modulates Kv2 subunit phosphorylation and clustering

Kv2 channels cluster at ER-PM junctions by interacting with VAP proteins via the PRC motif in their C-terminal domain (Johnson et al., 2018; Kirmiz et al., 2018a). Although KvS subunits lack a PRC motif, we find that heteromeric Kv2/Kv5.1 channels can cluster at ER-PM junctions on neuronal soma (Fig 7a). This indicates that clustering does not require PRC motifs in all four channel subunits, but it remains possible that Kv2/Kv5.1 channels have a diminished propensity to localize to and organize ER-PM junctions. To assess this, we compared the coefficient of variation (CV: SD/mean) of Kv2.1, Kv2.2 and Kv5.1 immunolabeling intensity in L2/3 neurons in cortex. The CV is used as a measure of non-uniformity of subcellular distribution, with clustered distributions having high CV values and uniform or diffuse signals having low CV values (Bishop et al., 2015; Kirmiz et al., 2018b; Kirmiz et al., 2018a). Indeed, we found that the CV value for Kv5.1 immunolabeling was lower than that for Kv2.1 and Kv2.2 immunolabeling in the same neurons (Fig 7b), suggesting that Kv5.1-containing channels are less clustered than the total Kv2 channel population.

**Fig 7.**
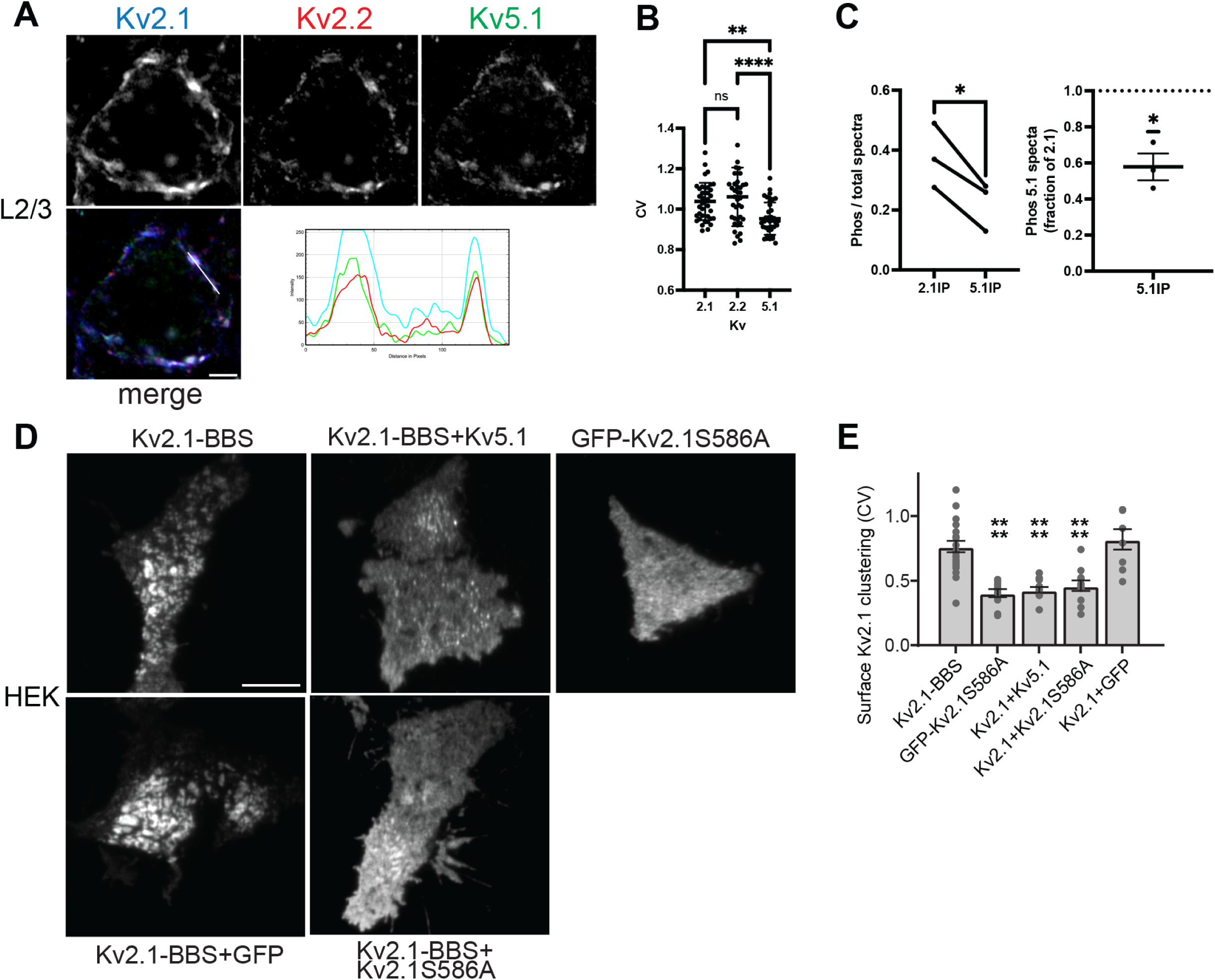
Kv5.1 reduces Kv2 channel clustering at ER-PM junctions. (A) Immunolabelling of L2/3 neurons shows that Kv5.1 (green) is co-clustered with Kv2.1 (blue) and Kv2.2 (red) on a neuronal soma, presumably at sites corresponding to ER-PM junctions. This is evident in the non-uniformity and coincidence of the three labels in the intensity profile (lower right). Scale bar = 2.5 µm. (B) Relative clustering of Kv2 and Kv5.1 subunits was compared by measuring the coefficient of variation of their labeling intensity (CV: SD/mean of pixel intensity). CV values for Kv5.1 were significantly lower than for either Kv2.1 or Kv2.2, suggesting that Kv5.1-containing channels are less clustered (one way ANOVA and Tukey’s multiple comparisons test, **** p<0.0001, ** p=0.0023, n=39 neurons, 2 WT brains). (C) Kv2.1 phosphorylation was assessed by proteomic analysis of Kv2.1 protein immunopurified from brain samples in IPs with Kv2.1 or Kv5.1 antibodies. Left panel. The fraction of spectra containing phosphorylated S/T residues on Kv2.1 was lower in Kv5.1-containing channels compared to total Kv2.1 channels (paired t test, * p=0.03, n=3). Right panel. Kv2.1 phosphorylation was decreased by ∼40% in Kv2/Kv5.1 heteromeric channels compared to all Kv2.1 channels (one sample t-test, * p=0.03, n=3). (D) Kv2.1BBS was expressed in HEK cells alone or together with Kv5.1 or Kv2.1S586A. Cell surface-expressed Kv2.1BBS was then labelled using AF647-Btx and imaged using TIRF. Kv2.1 is highly clustered when expressed alone but is more diffusely distributed when co-expressed with Kv5.1 or with Kv2.1 S586A. Scale bar = 10 µm. (E) Quantitation of CV values from transfected HEK cells. Co-expression of Kv5.1 and Kv2.1 S586A both resulted in a significant decrease in the CV value of surface Kv2.1BBS labeling, as compared to Kv2.1BBS expressed alone. Thus, Kv2.1/Kv5.1 and Kv2.1/Kv2.1S586A heteromeric channels both have a reduced propensity to cluster at ER-PM junctions (one way ANOVA and Tukey’s multiple comparisons test, **** p<0.0001); n = 21 [Kv2.1], 10 [S586A], 12 [Kv2.1 + Kv5.1], 11 [Kv2.1 + S586A], and 8 [Kv2.1 + GFP] cells).

Kv2 channel clustering has previously been shown to correlate with the phosphorylation status of Kv2 subunits (Misonou et al., 2004), which are phosphorylated at multiple S/T sites in their cytoplasmic C-termini (Park et al., 2006). Consequently, we performed a mass spectrometry-based analysis of Kv2.1 subunit phosphorylation in channels IPed from brain lysates using Kv2.1 or Kv5.1 antibodies. Interestingly, we found that the fraction of Kv2.1 spectra that contained phosphorylated S/T residues was ≈40% lower in Kv5.1-containing channels present in the Kv5.1 IPs as compared to the Kv2.1 present in IPs performed with anti-Kv2.1 antibodies (Fig 7c). As lower levels of Kv2.1 phosphorylation are typically associated with reduced channel clustering at ER-PM junctions, this supports our finding that Kv2/Kv5.1 heteromeric channels are less clustered than Kv2 channels in general.

Finally, we tested whether coexpression with Kv5.1 subunits also reduces clustering of Kv2 channels at ER/PM junctions in heterologous cells. In transfected HEK cells, we found that clustering of cell surface Kv2.1 was markedly reduced in cells co-expressing Kv5.1 compared to cells expressing Kv2.1 alone (Fig 7d). Indeed, the CV value for cells co-expressing Kv2.1 and Kv5.1 was significantly lower than for cells expressing Kv2.1 alone (Fig 7e). This result phenocopied that of cells co-expressing Kv2.1 and a non-clustering Kv2.1 mutant (S586A)(Kirmiz et al., 2018b), which would similarly reduce the number of functional PRC motifs in Kv2.1 channels. Thus, in heterologous cells, Kv2.1/Kv5.1 heteromeric channels cluster much less efficiently than Kv2.1 homomeric channels. Although more modest reductions in the clustering of Kv5.1-containing channels are observed in L2/3 neurons in brain, these findings indicate that Kv2/Kv5.1 heteromeric channels constitute a distinct subpopulation, which have a reduced ability to localize to and presumably to organize ER-PM junctions.

## Discussion

The Kv2 family is unusual amongst voltage-gated potassium channels in that it consists of only two alpha subunit family members (Kv2.1 and Kv2.2), each encoded by a single transcript without splice variants, limiting the structural and functional diversity of Kv2 homo- or hetero-tetrameric channels. We provide evidence here, however, that Kv2 channel subunits also co-assemble with the “electrically silent” or KvS subunit family, creating diverse Kv2 channel subtypes in brain.

Gene expression studies show that KvS subunit family members have restricted regional and cellular expression patterns in mouse brain, which partially overlap with that of Kv2.1 and Kv2.2 (e.g., Allen Brain Atlas, Mousebrain.org)(Bocksteins, 2016). For example, Kv5.1 (KCNF1) mRNA is broadly expressed in cortex (Drewe et al., 1992), Kv8.1 is expressed in cortex and hippocampus in a subset of excitatory neurons ((Castellano et al., 1997), Allen Brain Transcriptomics), Kv9.2 is expressed in hippocampus in CA1-3 and dentate gyrus ((Salinas et al., 1997a), Allen Brain Atlas), and Kv6.3 is expressed in hippocampus only in dentate gyrus (Allen Brain Atlas). Consistent with their differing mRNA expression patterns, our mass spectrometry analysis of Kv2 channel proteins identified multiple KvS subunits that varied significantly in their relative abundance. In native channels IPed with a Kv2.1 antibody, we found that spectral counts for Kv5.1 were ≈18% of that for Kv2.1, indicating that Kv2.1/Kv5.1 heteromers are somewhat common in brain. Other KvS subunits were detected at lower abundance in immunopurified Kv2.1 complexes (Fig 1a, Fig 2d). These counts may be an underestimate, as KvS subunits (∼50-55kD) are approximately half the size of Kv2.1 (∼100kD), which would likely result in fewer spectra being detected per KvS molecule compared to Kv2.1. Moreover, when taken in aggregate, the spectral counts for all five KvS subunits combined are ∼30% of that for Kv2.1, suggesting that a substantial subpopulation of native Kv2 channels in brain contain a KvS subunit.

Our proteomic and biochemical analyses reveal several additional key insights into Kv2 and Kv5.1 channel composition in brain. First, our experiments allow for some estimates on the frequency of Kv2.1/Kv2.2 heteromers in brain. Proteomic analysis of Kv2.1-containing channels identified Kv2.2 at ∼25% of the spectral abundance of Kv2.1. Similarly, IP and IB analysis showed that Kv2.1 IPs pull down ∼21% of total Kv2.2 subunit, and Kv2.2 IPs pull down ∼22% of total Kv2.1. Thus, we estimate that approximately 20% of Kv2.1 and Kv2.2 subunits occur as Kv2.1/2.2 heteromers. Second, immunoblotting of Kv2.1 and Kv2.2 IPs showed that more Kv5.1 is associated with Kv2.1 than Kv2.2 subunits, and mass spectrometry analysis of Kv5.1 IPs identified 2-fold higher spectral counts of Kv2.1 than Kv2.2. Thus, Kv2.1/Kv5.1 heteromers predominate over Kv5.1/Kv2.2 heteromers in brain. Third, while multiple KvS subunits were detected in Kv2.1 IPs, no additional KvS subunits were identified in Kv5.1 IPs. This indicates that Kv2/Kv5.1 channels typically do not contain more than one type of KvS subunit. Potentially, this could be due either to non-overlapping expression patterns of KvS subunits, molecular constraints that prohibit co-assembly of distinct KvS subunits, or only one KvS subunit being permitted in each tetrameric channel. Fourth, there is currently considerable debate over the stoichiometry of Kv2/KvS channels with evidence from heterologous cell experiments for Kv2 to KvS subunit ratios of either 2:2 (Moller et al., 2020) or 3:1 (Kerschensteiner et al., 2005; Pisupati et al., 2018). In Kv5.1 IPs from brain, we find that the ratio of spectral counts for Kv2 to Kv5.1 subunits is ∼3.5:1, and ∼2.3:1 when normalized for either the number of observed unique peptides or expected tryptic peptides per subunit. Although the subunit stoichiometry likely falls within this range, due to the semi-quantitative nature of spectral counting and differences in size of Kv2 and Kv5.1 subunits, we are unable to definitively resolve the predominant Kv2/KvS subunit stoichiometry from this data.

Additional evidence that Kv5.1 and Kv2 subunits form heteromeric channels is that Kv5.1 protein levels are decreased in Kv2.1 KO mice and even more so in Kv2.1/2.2 dKO mice. Similarly, Kv5.1 immunolabeling was greatly reduced or often undetectable in Kv2.1 KO neurons, and any residual Kv5.1 co-localized with remaining Kv2.2 subunit. Thus, similar to the AMIGO-1 auxiliary subunit (Bishop et al., 2018), Kv5.1 protein expression in brain neurons depends on its obligate Kv2 subunit partners and presumably their co-assembly into heteromeric channels, whereas unassembled Kv5.1 subunits are likely unstable and rapidly degraded.

At the regional and cellular level, our immunolabeling of mouse brain sections for Kv5.1 showed that it is predominantly expressed in neocortex and differentially expressed across cortical layers. Kv5.1 immunolabeling is highest in L2-3, where it is robustly present in a large subset of Kv2 positive neurons, but it is also detected in smaller subsets of Kv2 positive neurons in deeper cortical layers (L4-6). This pattern of Kv5.1 immunolabeling closely resembles its mRNA expression in cortex, as shown by both in situ hybridization and transcriptomics studies. In individual neurons, we found that Kv5.1 immunolabeling colocalizes precisely with Kv2.1 and Kv2.2, providing further evidence that they form heteromeric channels. Interestingly, however, the relative levels of Kv2.1, Kv2.2 and Kv5.1 immunolabeling vary widely between neurons even in the same cortical layer, suggesting that there is remarkable heterogeneity in Kv2 channel composition between neurons. Thus, Kv5.1 likely contributes to Kv2 channel function to varying degrees in different neurons.

The co-assembly of Kv5.1 with Kv2 subunits to form heteromeric channels has the potential to modify multiple aspects of Kv2 channel function. Previous studies have shown that Kv2 channels regulate neuronal excitability in a complex manner that depends on patterns of excitation (Liu and Bean, 2014; Speca et al., 2014; Honigsperger et al., 2017; Romer et al., 2019), and that KvS subunits modify the biophysical properties of Kv2 channels in a subunit-specific manner (Bocksteins, 2016). Consistent with prior findings (Kramer et al., 1998), we find that Kv5.1 shifts the voltage-dependence of Kv2 channel activation to more positive membrane potentials and slows channel deactivation. Thus, the presence of Kv2/Kv5.1 heteromeric channels as well as Kv2 homomeric channels in select cortical neurons may well influence the excitability and firing patterns of those neurons. Moreover, Kv2.1/Kv5.1 heteromers have altered sensitivity to inhibition by RY785, broadening the impact of KvS subunits on Kv2 channel function and impacting interpretation of experiments employing this selective inhibitor of Kv2 channels in native cells.

Notably, KvS subunits also modify the pharmacological response of Kv2/KvS heteromers to another pore blocker, 4-aminopyridine (Stas et al., 2015). Another key function of Kv2 channels is to organize ER-PM junctions on neuronal somata and proximal dendrites (Johnson et al., 2019; Vierra and Trimmer, 2022). This structural role is mediated by phosphorylation-dependent binding of the Kv2.1 PRC domain to ER-localized VAP proteins (Johnson et al., 2018; Kirmiz et al., 2018a). Notably, all KvS subunits lack a PRC domain, and consistent with this, we found that Kv5.1 co-expression significantly reduces Kv2 channel clustering in heterologous cells. Similarly, we found that Kv5.1 is slightly less clustered than either Kv2.1 and Kv2.2 in L2/3 cortical neurons in brain. Moreover, Kv2.1 subunits in Kv5.1-containing channels were ∼40% less phosphorylated compared to Kv2.1 channels in total, consistent with a less-clustered distribution (Misonou et al., 2004). Taken together, these findings suggest that the decreased number of PRC domains in Kv2/Kv5.1 heteromeric channels reduces their propensity to cluster at ER-PM junctions, likely by changing the avidity of VAP interactions. Finally, Kv2 channels help assemble a macromolecular complex at ER-PM junctions that includes LTCCs, Ca^2+^ signaling and scaffolding proteins (Vierra et al., 2019), lipid handling proteins (Kirmiz et al., 2019), and PKA signaling proteins (Vierra et al., 2023), thereby creating a specialized Ca^2+^, lipid and PKA signaling microdomain. As many of these interactions are mediated - directly or indirectly - by the large cytoplasmic C-termini of Kv2 subunits, replacement of Kv2 with Kv5.1 in channel tetramers could impact the channel interactome and composition of the associated ER-PM junctions. Most obviously, Kv2/Kv5.1 channels could potentially interact with fewer of these junctional components, reducing their localization at ER-PM junctions, but it is also possible that Kv5.1 might recruit select proteins though direct binding. Thus, important questions for future studies are whether Kv5.1 and other KvS subunits modulate the ability of Kv2 channels to localize to and organize ER-PM junctions, and if they impact Ca^2+^ and lipid signaling functions at these specialized junctional complexes in brain neurons.

## Acknowledgements

This work was supported by NIH research grants R21NS123417 to MF, R03TR004200 to MF and JS, and R01NS114210 and R21NS101648 to JST. We thank members of the Ferns, Sack and Trimmer laboratories for their support and useful discussions of the data.

