## Supplemental Figure 1 for "Electrically silent KvS subunits associate with native Kv2 channels in brain and impact diverse properties of channel function"

### Suppl. Fig. 1

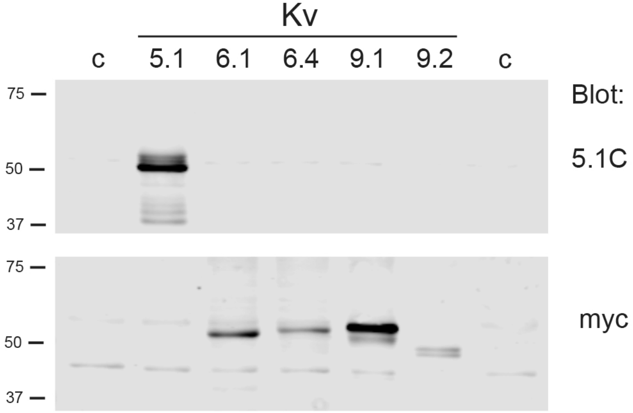


**S1. Specificity of Kv5.1C antibody**.
HEK cells expressing Kv5.1 and myc tagged Kv6.1, Kv6.4, Kv9.1 and Kv9.2 were solubilized, size fractionated on SDS gels and immunoblotted with 5.1C pAb (top) and myc mAb (bottom). Affinity-purified 5.1C pAb detected Kv5.1 but not other KvS subunit proteins. Numbers to the left are molecular weights standards in kD.
